# Extracellular succinate induces spatially organized biofilm formation in *Clostridioides difficile*

**DOI:** 10.1101/2023.01.02.522461

**Authors:** Emile Auria, Julien Deschamps, Romain Briandet, Bruno Dupuy

## Abstract

*Clostridioides difficile* is the major cause of nosocomial diarrhea, which are associated with gut microbiome dysbiosis. Biofilms of *C. difficile* have been progressively linked to the pathogenesis of this bacterium and the recurrences of its infections. Though the number of conditions in which *C. difficile* biofilms are being produced is increasing, little is known about how and when biofilms are formed in the gut. Here we report that succinate, a metabolite abundantly produced by the dysbiotic gut microbiota, induces *in vitro* biofilm formation of *C. difficile* strains. We characterized the morphology and spatial composition of succinate-induced biofilms, and compared to non-induced or deoxycholate-induced biofilms, biofilms induced by succinate are significantly thicker, structurally more complex, and poorer in proteins and exopolysaccharides (EPS). We then applied transcriptomics and genetics to characterize the early stages of succinate-induced biofilm formation and we showed that succinate-induced biofilm results from major metabolic shifts and cell-wall composition changes. Similar to deoxycholate-induced biofilms, biofilms induced by succinate depend on the presence of a rapidly metabolized sugar. Finally, although succinate can be consumed by the bacteria, we found that the extracellular succinate is in fact responsible for the induction of biofilm formation through complex regulation involving global metabolic regulators and the osmotic stress response. In the context of human gut dysbiosis, succinate can limit bacterial infections through the control of innate immune responses. Collectively, our results suggest that succinate is an intestinal signal which can drive the biofilm formation and persistence of *C. difficile* in the gut and increase the risk of relapse.

## Introduction

Dysbiosis caused either by inflammatory bowel disease or antibiotic intake can develop serious complications, the most common one being *Clostridioides difficile* infections (CDI), the leading cause of nosocomial diarrhea and colitis in industrialized countries. *C. difficile* is an anaerobe spore-forming bacterium known to be an opportunistic gut pathogen and a rising public health concern due to its increasing resistance to the antibiotics used to treat the infections (1). In addition, CDI present a high recurrence rate, between 15% and 35% of all CDI cases. In 40% of recurrent cases, patients are infected with the same strain that caused the initial infection, suggesting *C. difficile* is able to persist in the gastrointestinal tract (2). Persistence can partly be attributed to *C. difficile* spores being formed and staying in the gut either in the microbial communities (3) or in the epithelial cells (4). In addition, it has also been suggested that *C. difficile* can persist in the gut in the long run through biofilm formation, allowing for more frequent relapses (3). Biofilms are microbial communities encased in an autoproduced polymeric matrix attached to a substrate, allowing them to better colonize and survive in a given environment (5). *C. difficile* can form biofilms *in vitro* in regular culture medium in several days (6–8) or faster when induced by specific molecules (9,10). Moreover, biofilm-like structures have also been observed in mono-associated mouse models (11), confirming that *C. difficile* biofilm formation can indeed occur in the gut.

Several gut-relevant signals have been shown to promote biofilm formation, including microbiota-produced metabolites such as short-chain fatty acids (12,13). Inorganic molecules like iron and phosphate can induce biofilms, as well as the presence of subinhibitory concentrations of antimicrobial molecules and host-derived signals such as antibiotics and bile salts, respectively (14). In *C. difficile*, the induction of biofilm formation was observed in response of sub-inhibitory concentrations of vancomycin (15) and metronidazole (9), the most commonly used treatments against mild CDIs, as well as in response to the secondary bile salt deoxycholate (10). However, despite the identification of regulatory mechanisms and growth conditions controlling *C. difficile* biofilm formation (16), very little is known about specific signals or inducers involved in biofilm formation.

Among the microbiota-produced metabolites that prevent bacterial infections, succinate has been shown to be potentially involved in tuning the immune response towards bacterial infection control (17). Succinate is also a key metabolite in the tri-carboxylic acid or Krebs cycle, as well as a key intermediate in propionate fermentation in bacteria (18). In both cases, its role resides in energy metabolism through the reduction of FAD and NAD+ cofactors.

Recently, it has been shown that succinate is a critical metabolite that promotes high levels of inflammatory cytokine IL-1β by inhibiting the negative regulator of hypoxia-induced factor 1α (HIF-1α) (19), and modulates distinct phases of the innate immune response required to clear bacterial infections. Both innate cellular immunity and innate humoral immunity that constitute the innate immune response are interchangeably regulated by succinate and inosine, according to their levels in the gut in a time-dependent manner (17).

As a product of microbial metabolism, the presence of succinate in the gut is mostly due to specific bacterial species, including several Bacteroides and Negativicutes that are present in the commensal colonic microbiota (20). In addition, other gut bacterial species such as *Phascolarctobacterium succinatutens* (21) or *C. difficile* (22) thrive on succinate when they are present in the gut, participating in maintaining its concentration low in the gut. Without these succinate consumers, the production of succinate by gut commensals could be detrimental to the host as higher concentrations of succinate are toxic for colonocytes and other gut cells and bacteria (23,24). Higher concentrations of luminal succinate have been measured both in humans and animal models in cases of gut microbiota dysbiosis (22,25,26). Those can be caused by a variety of situations such as inflammatory bowel diseases like Crohn’s disease or antibiotic treatments (27,28).

Although *C. difficile* encounters succinate in relatively high concentrations during colonization of the dysbiotic intestine, little is known of the impact of this metabolite on the bacterium besides its ability to be used for its growth (22). Poquet and collaborators (2018) reported that in biofilm cells the reductive utilization of succinate is active, and genes involved in its uptake and metabolism are up-regulated (29). Here we report that succinate induces biofilm formation in various *C. difficile* strains at physiologically relevant concentrations. We demonstrated that extracellular succinate, and not the one used by the bacteria, is the key metabolite that triggers biofilm formation. We then identified the main factors that could contribute to the succinate-induced biofilms by performing transcriptomic experiments and using a set of mutants, and compared the biofilm structure of succinate and DCA-induced biofilms by CLSM experiments. Our results highlight the diversity of biofilm formation in *C. difficile* depending on the inducers and its probable importance in the process of gut colonization.

## Materials and methods

### Bacterial strains and culture conditions

The bacterial strains and plasmids used in this study are listed in Table S1. All *C. difficile* strains were grown in anaerobic conditions (5% H2, 5% CO2, 90% N2) in TY medium (30g/L tryptone, 20g/L yeast extract) or in biofilm growth BHISG medium (BHI with 0.5% (w/v) yeast extract, 0.01 mg/mL cysteine and 100mM glucose). *C. difficile* semi-defined biofilm growth medium (BM) was also used for biofilm assays with the following composition (30): Oxoid casein hydrolysate (10 mg/mL), L-Tryptophane (0.5 mg/mL), L-Cysteine (0.01 mg/mL), L-Leucine (0.0033 mg/mL), L-Isoleucine (0.0033 mg/mL), L-Valine (0.0033 mg/mL), Na2HPO4 (5 mg/mL), NaHCO3 (5 mg/mL), KH2PO4 (0.9 mg/mL) NaCl (0.9 mg/mL), (NH4)2SO4 (0.04 mg/mL), CaCl2·2H2O (0.026 mg/mL), MgCl2·6H2O (0.02 mg/mL), MnCl2·4H2O (0.01 mg/mL), CoCl2·6 H2O (0.001 mg/mL) FeSO4·7 H2O (0.004 mg/mL) D-biotine (0.001 mg/mL), calcium-D-panthothenate (0.001 mg/mL) and pyridoxine (0.1 μg/mL). This medium was supplemented with glucose or other sugar sources when necessary. BM supplemented with 100mM of glucose is noted BMG.

### Growth curves

Cell growth was monitored by manually measuring the OD_600nm_ from 10mL cultures in BM medium supplemented with succinate (80mM) and/or glucose (100mM) or cellobiose (100mM).

### Gene deletion in *C. difficile*

Gene deletion in *C. difficile* was performed as described in Peltier *et al*. (Peltier et al 2020). Regions upstream and downstream of the genes of interest were amplified by PCR using primer pairs described in Table S1. PCR fragments and linearized pDIA6754 (31) were mixed and assembled using Gibson Assembly (NEB, France) and transformed by heat shock in *E. coli* NEB 10β strain. The plasmid constructions were verified by sequencing and plasmids with the right sequences were transformed in *E. coli* HB101 (RP4). The resulting strains were used as donors in a conjugation assay with the relevant *C. difficile* strains. Deletion mutants were then obtained using a counter-selection as described in Peltier *et al*. (2020).

### Biofilm assays

Overnight cultures of *C. difficile* grown in TY medium with appropriate antibiotics were diluted to 1/100 into fresh BHISG or BMG containing supplements (sodium or potassium succinate 20-120mM, sodium butyrate 20-200mM, DCA 240μM) when needed. Depending on the assay, the diluted overnight cultures were aliquoted either with 1mL per well in 24-well plates (polystyrene tissue culture-treated plates, Costar, USA) or with 200μL per well in 96-well plates (polystyrene black tissue-culture-treated plates, Greiner Bio One, Austria). The plates were incubated at 37°C in an anaerobic environment for 24h or 48h. 24-well plates were used for RNA-isolation. For biofilm assays in 96-well-plates, two different treatments were performed depending on the assay. Plates used for biofilm biomass quantification were treated in a similar manner as the established method (Dubois et al 2019) with an additional step. Briefly the medium was removed by inverting the plate and wells were washed once by pipetting gently 200μL of phosphate-buffer saline (PBS) which was removed by inverting the plate. 200μL of fixing solution (75% ethanol, 25% acetic acid) were thereafter added to the wells and the plate was incubated at room temperature for 20min. The fixing solution was removed by inverting the plate and the wells were air dried for 1h at room temperature. Then 200μL of a crystal violet solution (0.2%) were added to the wells and incubated for 10min at room temperature before removing the staining solution by inverting the plate. The wells were gently washed twice with 200μL PBS then 200μL of 70% ethanol were added to the wells. The absorbance, corresponding to biofilm biomass was measured at a wavelength of 600nm with a plate reader. The plates used for CLSM were treated differently. First, the spent medium was carefully removed by pipetting then 200μL PBS supplemented with 4% of paraformaldehyde (PFA) were added. Plates were incubated for an hour at room temperature and the media was carefully removed by pipetting before adding PBS for 48h at 4°C. Dyes were then directly added to the wells for biofilm matrix imaging. Plates used for live-dead microscopy were untreated before the addition of the dyes. In all assays, a sterile medium was used as the negative and blank controls for the assays.

### RNA isolation

Cells were grown in 24-well plates and 10 wells per plate were used to produce one replicate for one condition. For RNA isolated from biofilms, the supernatant was removed by inverting the plate and the biofilms were carefully washed twice and resuspended in 3 mL of PBS. In other conditions, the whole bacterial population was collected, and cells were harvested by centrifugation (10 min, 5000 rpm, 4°C) and resuspended in 1 ml of PBS. Cell suspensions in PBS were finally centrifuged (10 min, 5000 rpm, 4°C) and the pellets were frozen at -80°C until extraction of total RNA performed as described in Saujet *et al* (2011) (32).

### Whole transcriptome sequencing and analysis

Transcriptome analysis for each condition was performed with 3 independent RNA preparations. Libraries were constructed using an Illumina Stranded Total RNA Prep Ligation with RibZero Plus (Illumina, USA) according to the supplier’s recommendations. RNA sequencing was performed on the Illumina NextSeq 2000 platform using 67 bases for a target of 10M reads per sample.

### Confocal Laser Scanning Microscopy (CLSM)

Biofilms grown in 96-well plates (Microclear, Greiner Bio-one, France) in BHISG supplemented with either DCA (240μM) or succinate (120mM) were obtained and fixed with PFA as described above. The final concentrations of the dyes in the wells were as follows i.e., 20μM SYTO9 (Life Technologies, USA); 20μM SYTO61 (Life Technologies); 20μM propidium iodide; 100nM TOTO-1 (Invitrogen, USA); FilmTracer SYPRO Ruby (manufacturer’s guidelines) (Invitrogen) and 50μg/mL calcofluor white (Merck, USA). Dyes were incubated for 30min before CLSM imaging/analysis. Z stacks of horizontal plane images were acquired in 1 μm steps using a Leica SP8 AOBS inverted laser scanning microscope (CLSM, LEICA Microsystems, Wetzlar, Germany) at the INRAE MIMA2 platform (33). At least two stacks of images were acquired randomly on three independent samples at 800 Hz with a x63 water objective (N.A.=1.2). Fluorophores were excited, then their emissions were captured as prescribed by the manufacturer.

### Analysis of CLSM biofilm images

Z-stacks from the CLSM experiments were analyzed with the BiofilmQ software (Hartmann et al 2021) to extract quantitative geometric descriptors of biofilms structures. Images were all treated with the same process in each fluorescence channel. First, the images were denoised by convolution (dxy=5 and dz=3), then they were segmented into two classes with an OTSU thresholding method with a sensitivity of 2. The detected signal was then declumped in 3.68μm cubes and small objects were removed with a threshold of (0.5μm^3^) to clean the remaining noise. Exported data were analyzed in the software Paraview v5.11 to generate biofilm 3D projections and in R (ggplot2 library) to generate quantitative graphs.

### Succinate quantification

Succinate was quantified in the spent media of 24h BMG cultures of *C. difficile* using the succinate colorimetric assay kit (Merck) according to the manufacturer’s guidelines. Results take evaporation into account and were normalized using uninoculated media.

### Statistical analysis

Biofilm biomass assays using crystal violet coloration were analyzed with the following tests when appropriate: i) a Brown-Forsythe and Welch ANOVA test followed by an unpaired t test with Welch correction or followed by Dunnett’s multiple comparisons test, ii) an ordinary one-way ANOVA test followed by Dunnett’s multiple comparisons test or followed by an uncorrected Fisher’s LSD and iii) an unpaired t-test. CLSM quantitative data were analyzed with Wilcoxon tests, while succinate quantification was analyzed with a paired t-test.

### Data Availability

RNA-Seq data generated in this study are available in the NCBI-GEO with accession no XXXXXXXX.

## Results

### Succinate induces biofilm formation in *C. difficile*

During dysbiosis, succinate is present in the gut at a mean concentration equivalent to 15mM to 60mM of succinate, depending on the density of the caecal content (22). Therefore, we performed biofilm assays in the biofilm growth medium BMG supplemented with concentrations of succinate ranging from 20mM to 80mM (Figure 1a). After 48h of growth, we observed biofilm formation in response to succinate in a dose-dependent manner. Since we used di-sodium succinate as the source of succinate, we performed two control experiments to verify the specificity of succinate as an inducer. First, we replaced succinate with butyrate in similar concentrations and no biofilm was formed (Figure 1b). Then, we used the counterion potassium with di-potassium succinate in identical concentrations and biofilm was produced in a similar quantity than with di-sodium succinate (Figure 1c). Thus, we confirmed that succinate can induce biofilm formation in *C. difficile* 630Δ*erm* strain.

**Figure 1:**
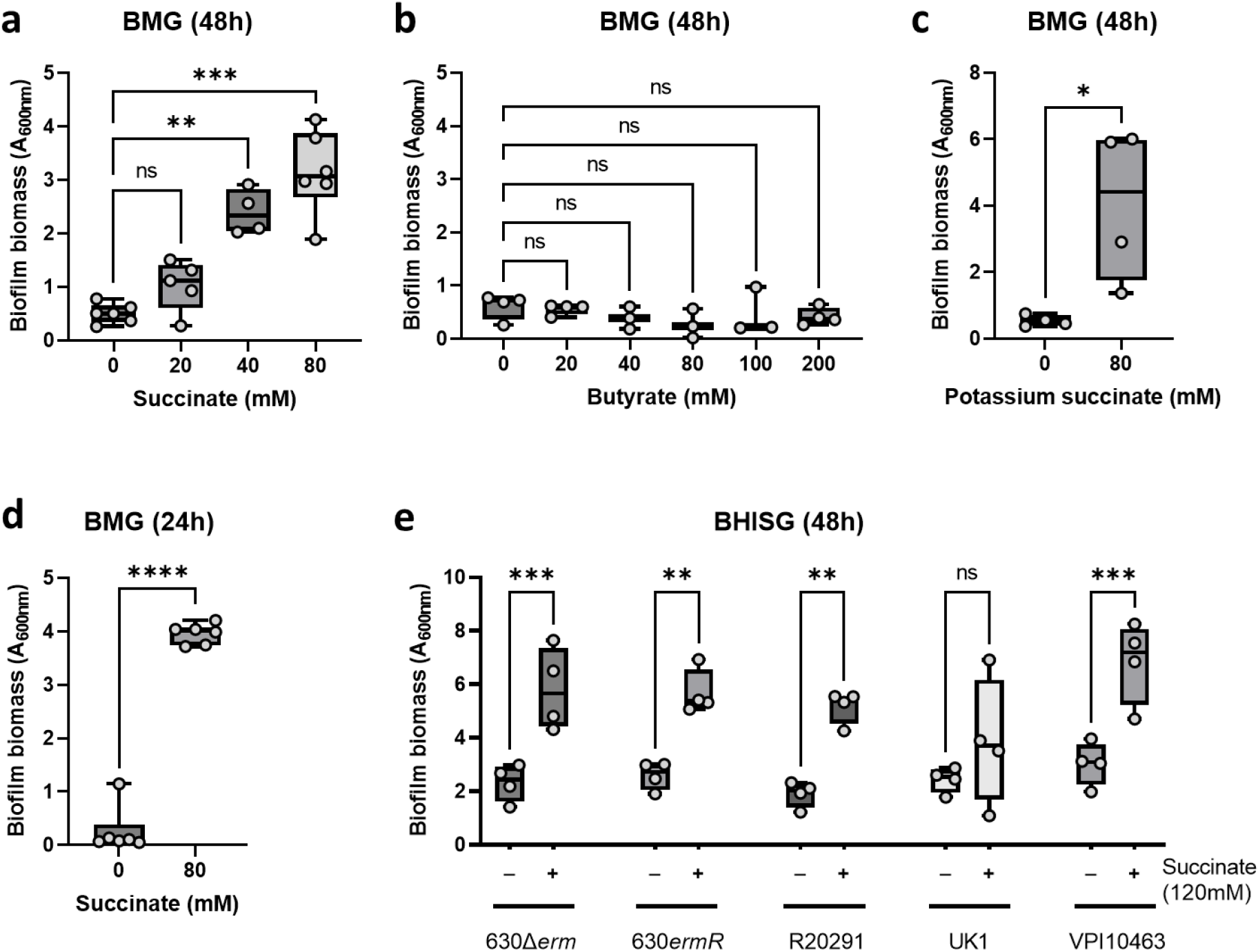
Succinate induces biofilm formation in *C. difficile*. Biofilm formation of the 630Δ*erm* strain was assayed in BMG medium supplemented with: **a**. sodium succinate at 48h, **b**. sodium butyrate at 48h, **c**. potassium succinate at 48h and **d**. sodium succinate at 24h. **e**. Biofilm formation of clinical strains was assayed in BHISG or BHISG supplemented with 120mM of sodium succinate. Each data point represents an independent biological replicate that is the mean of four technical replicates. Statistical analyses performed are a) and e) a Brown-Forsythe and Welch ANOVA test followed by an unpaired t-test with Welch’s correction, b) an ordinary one-way ANOVA test followed by Dunnett’s multiple comparisons test, c) and d) an unpaired t-test and (ns: not significant; *: p<0.05; **: p<0.01; ***: p<0.001).

As the biofilm biomass formed in response to succinate was plentiful at 48h of incubation, we wondered whether biofilm could be induced earlier. We found that contrary to other inducers such as deoxycholate (10), succinate induces biofilm formation as early as 24h of incubation (Figure 1d) and reaches the same biomass levels as those of 48h of incubation. Finally, to ensure that biofilm-induction by succinate was not strain-specific, the effect of succinate was tested on several clinical isolates such as R20291, UK1 and VPI10463 in BHISG medium (Figure 1e). This medium was used instead of BMG as several of the tested strains did not grow properly in BMG medium, which was optimised for the 630Δ*erm* strain (data not shown). In addition, the concentrations of succinate used in this medium are higher as well, as succinate-induced biofilms do not occur below 120mM of succinate in BHISG (data not shown). We showed that succinate can induce biofilm formation in all the tested strains in similar quantities as in the 630Δ*erm* strain (Figure 1e). Conjointly, the data shows that succinate strongly and specifically induces biofilm formation in *C. difficile* and that induction is not strain-dependent.

### Metabolic requirements of succinate-dependent biofilm induction

Since glucose is known to widely regulate *C. difficile* transcription (34) and the biofilm growth media used contained 100mM of glucose, we tested whether induction of biofilm by succinate depended on the presence of glucose. When we performed the biofilm assay in the presence of succinate with decreasing amounts of glucose in the medium, we found that no biofilm was formed at lower concentrations of glucose, and succinate-induced biofilm formation needs a high concentration of glucose to take place (Figure 2a). Interestingly, the induction does not seem to be glucose dose-dependent, rather, 80mM seems to be the lower threshold at which biofilm formation can occur in response to succinate. As glucose is not the only available sugar source in the gut, we tested whether other sugars could replace glucose in the succinate-induced biofilm formation. As shown in Figure 2b, similar concentrations (100mM) of mannose, N-acetyl glucosamine (NAG) can replace glucose in the succinate-induced biofilm formation, whereas cellobiose and galactose cannot. The replacement of glucose by mannose in biofilm formation was already observed by Piotrowski and collaborators (2019) when they used mannose in similar concentrations as glucose as a biofilm enhancer of the *C. difficile* 630 strain (35). We confirmed the essential role of PTS transported sugars such as glucose or mannose in succinate-induced biofilm formation by testing in biofilm assays a *ptsI* gene mutant, encoding an enzyme involved in the transport of PTS sugars. Succinate-induced biofilm formation was completely abolished in the *ΔptsI* strain (Figure 4a). We noted that although cellobiose cannot replace glucose in the succinate-induced biofilm formation, *C. difficile* can use cellobiose as a carbon source (Figure 2d). Moreover, cellobiose is metabolically close to glucose as a glucose dimer (36). In order to assess the specificity of glucose in the succinate-dependent biofilm induction, we progressively replaced glucose by cellobiose in the biofilm growth medium while keeping a total concentration in carbohydrates of 100mM (Figure 2c). The results show that biofilms are induced by succinate at a threshold similar to that corresponding to 100mM glucose concentration from 70mM glucose supplemented with 30mM cellobiose in the medium. Since cellobiose is metabolized by *C. difficile* more slowly than glucose (Figure 2d), it seems that a rapidly metabolized sugar is needed for succinate-dependent biofilm induction, which can be replaced later by a slowly metabolized sugar (Figure 2d).

**Figure 2:**
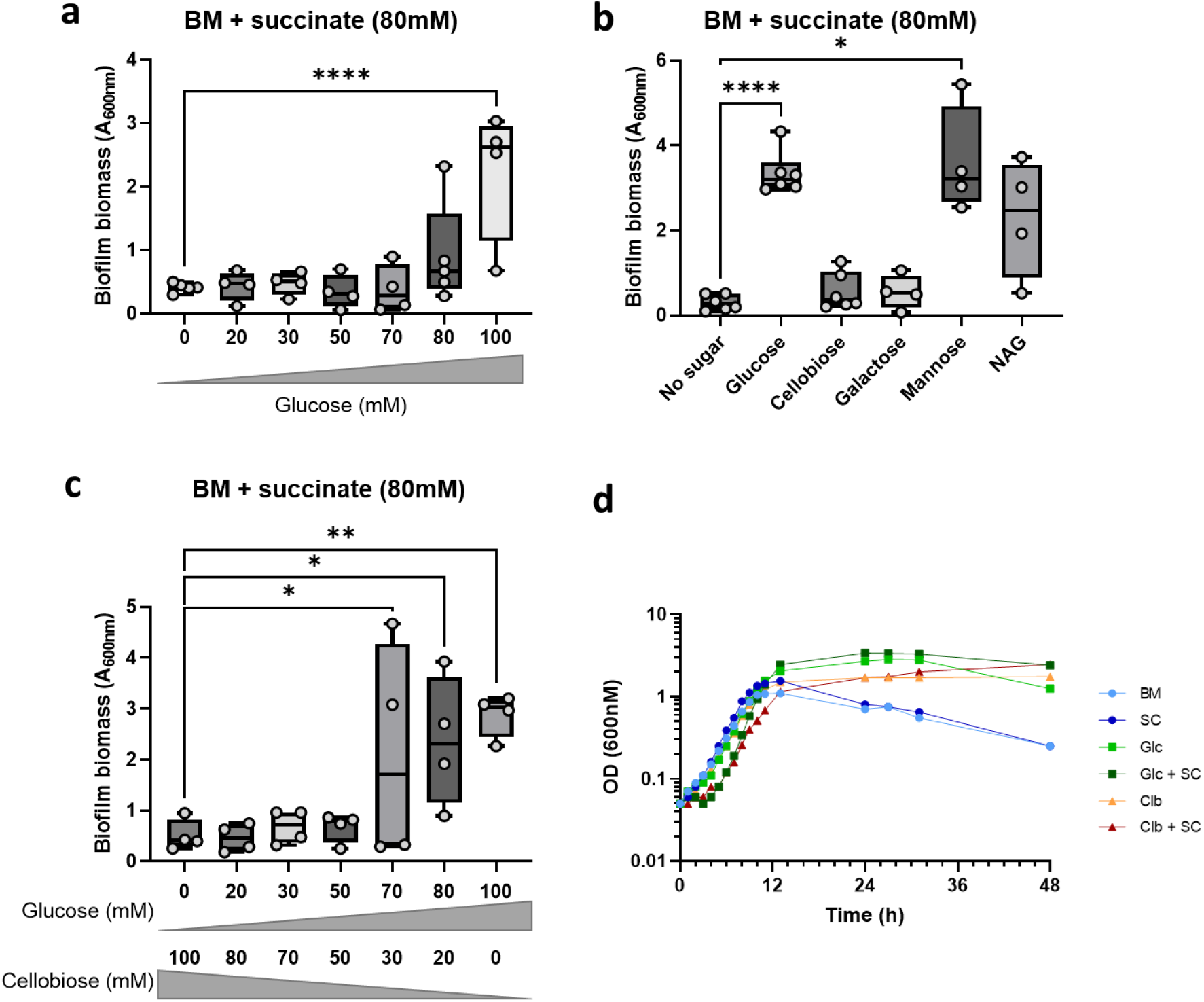
Importance of the sugar in succinate-induced biofilms. Biofilm formation of the 630Δ*erm* strain was assayed in BM medium supplemented with succinate (80mM) and a variety of sugars such as **a**. glucose at different concentrations, **b**. different sugars (cellobiose, galactose, mannose, NAG, xylose) at 100mM each compared to glucose and **c**. a mix of glucose and cellobiose at 100mM final concentration. **d**. Growth curves of the 630Δ*erm* strain in BM medium supplemented with succinate (80mM), glucose (100mM) or cellobiose (100mM) and succinate + glucose or succinate + cellobiose. For the biofilm assays each data point represents an independent biological replicate that is the mean of four technical replicates. The growth curves are representative of three independent biological replicates. SC: succinate (80mM); Glc: glucose (100mM); Clb: cellobiose (100mM). Statistical analyses performed are a) a Brown-Forsythe and Welch ANOVA test followed by an unpaired t-test with Welch correction, b) and c) an ordinary one-way ANOVA test followed by a Fisher’s LSD test (*: p<0.05; **: p<0.01; ***: p<0.0001).

### Transcriptomic characterization of the succinate induction of biofilm

To identify key events leading to biofilm formation induced by succinate, we performed a comparative transcriptomic analysis on the *C. difficile* 630Δ*erm* strain grown in BMG supplemented or not with 80mM of succinate and incubated either at 14h or 24h. No biofilm formation was observed at 14h growth, while it was at 24h. Comparing the differences in the regulated genes between 14h and 24h of incubation allowed the identification of specific common regulations occurring in succinate-induced biofilms. Most of them may refer to the physiological state of the bacteria at the onset of biofilm formation. In both transcriptomes, only genes with a fold change of >2 or <-2 and a significant p value (p<0.05) were considered. A total of 641 and 2137 genes were differentially regulated in the presence of succinate at 14h and 24h of incubation, respectively (Figure 3). Many cell function-associated genes were regulated during biofilm formation, and we decided to focus our analysis on some of them, known to contribute to biofilm formation (Table S2). At 14h of incubation, while biofilms are not yet formed, the regulation of genes encoding enzymes associated with metabolism had already switched from glycolysis and the Wood-Ljungdahl pathway to succinate and formate fermentation. Furthermore, up-regulated genes involved in Stickland fermentations are directed towards the utilization of proline and aromatic amino-acids, whereas glycine and leucine pathways are down-regulated. Genes of several amino acid biosynthesis such as histidine, isoleucine/valine, methionine and aromatic amino acids, as well as spermidine, ornithine, arginine and asparagine are up-regulated, while those of glutamate and glycine are down-regulated. Shifts also occur for some membrane transporters, with the down-regulation of mannose/fructose and mannitol transporters and the up-regulation of methionine and spermidine transporters. Interestingly, iron seems to be imported during the first stage of growth since the genes encoding its transport are globally upregulated, contrary to those involved in molybdenum and cobalt transport. The cell wall properties are probably modified during growth in the presence of succinate since genes involved in D-alanylation of the teichoic acids, which decrease the negative charge of the cell surface and increase its hydrophobicity, are up-regulated. Among known regulators whose encoding genes are up-regulated, important transcriptional regulators mainly associated with metabolism and phase transition, seem to be involved in the onset of the succinate induction of biofilms such as CodY and the SinRR’ system, known in other species such as *B. subtilis* to control biofilm formation (37), and the CsfT regulator that belongs to the extracytoplasmic function (ECF) sigma factors, which sense and respond to extracellular signals (38).

**Figure 3:**
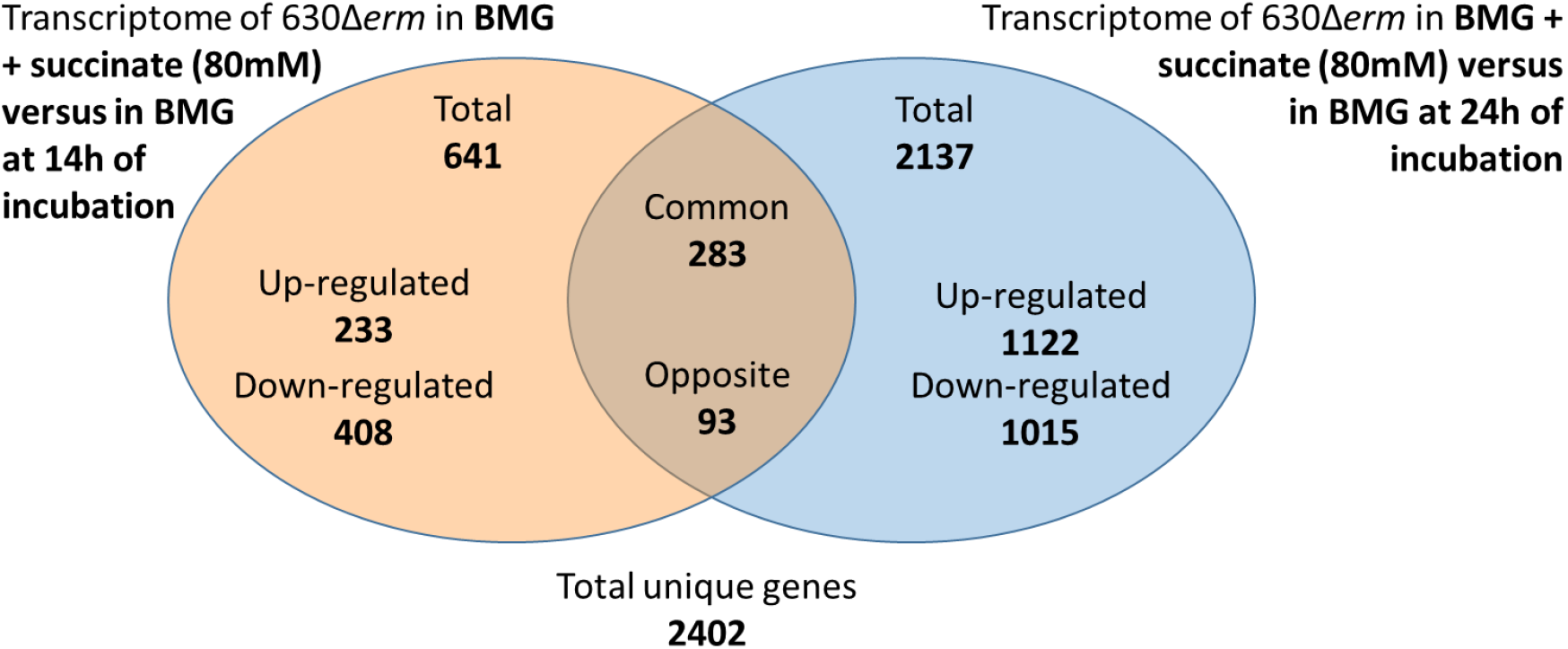
Differences in gene expression in the transcriptomic experiments. Venn diagram of the genes differentially regulated in the two transcriptomic experiments of *C. difficile* 630Δ*erm* strain grown in BMG medium supplemented or not with 80mM of succinate performed at 14h or 24h incubation.

At 24h, more significant transcriptional shifts occurred following those already observed at 14h of incubation in the presence of succinate, including major changes in metabolism (Table S2). Indeed, genes involved in formate use and Stickland fermentations are down-regulated, as are those involved in fatty acid catabolism. The genes associated with carbon catabolism pathways are still down-regulated at 24h and it seems that *C. difficile* gets its chemical energy from succinate fermentation. While the genes involved in glycolysis are down-regulated, several genes of membrane transporters of sugars such as those of fructose, maltose, glucose and other beta-glucosides are up-regulated. The latter carbohydrates could be used for peptidoglycan synthesis as a number of genes involved in these reactions are up-regulated at 24hrs. In addition, genes of several amino acid biosynthesis and transporters are up-regulated. This concerns the synthesis of thiamine, cysteine, ornithine, spermidine, and arginine, as well as the transport of glutamate and aspartate, two critical amino acids associated with multiple metabolic pathways, including protein synthesis and nucleotide metabolism. This is consistent with the up-regulation of *mreBCD* along with *ftsZ, ftsH* and *scpAB*, involved in cell shape determination and cell division, respectively. Most sporulation genes are still down-regulated and motility is significantly reduced since flagellar and pili genes are strongly down-regulated. This is consistent with the down-regulation of the regulator genes encoding *sigD* and *sigH* involved in the control of the expression of the flagella and the onset of sporulation, respectively. The metabolic rearrangements observed at 24hrs can be associated with the down-regulation of *rpoN* involved in the expression of the Stickland reaction genes and the up-regulation of *rex* and *codY* known to control genes of the carbon and nitrogen flux. Interestingly, both the SinRR’ system and the SigH sigma factor are up-and down-regulated, respectively, at 24h of incubation, indicating the probable importance of these regulators in the control of succinate-dependent biofilm induction. We know that SigH and Spo0A contribute to *C. difficile*’s metabolic adaptation (32,39). Moreover, SinR modulates Spo0A expression (40). Since *spo0A* inactivation decreased biofilm formation (Dubois et al, 2019, Fig 4d), we tested the effect of SigH and SinR on biofilm formation. Inactivation of *sinRR’* resulted in highly increase biofilm-formation whereas inactivation of *sigH* had no effect (Fig 4d). In addition, contrary to the 14h incubation, *CD1383* (a *busR*-like regulator, (41)) and *agrD* genes are upregulated at 24h, suggesting that succinate induction may be associated with the osmotic stress response and/or quorum sensing (QS). The role of QS is supported by the decrease in biofilm formation in the *luxS* mutant, but not by the *agrBD* mutant whose biofilm is elevated for reasons that are not understood (Figure 4d).

**Figure 4:**
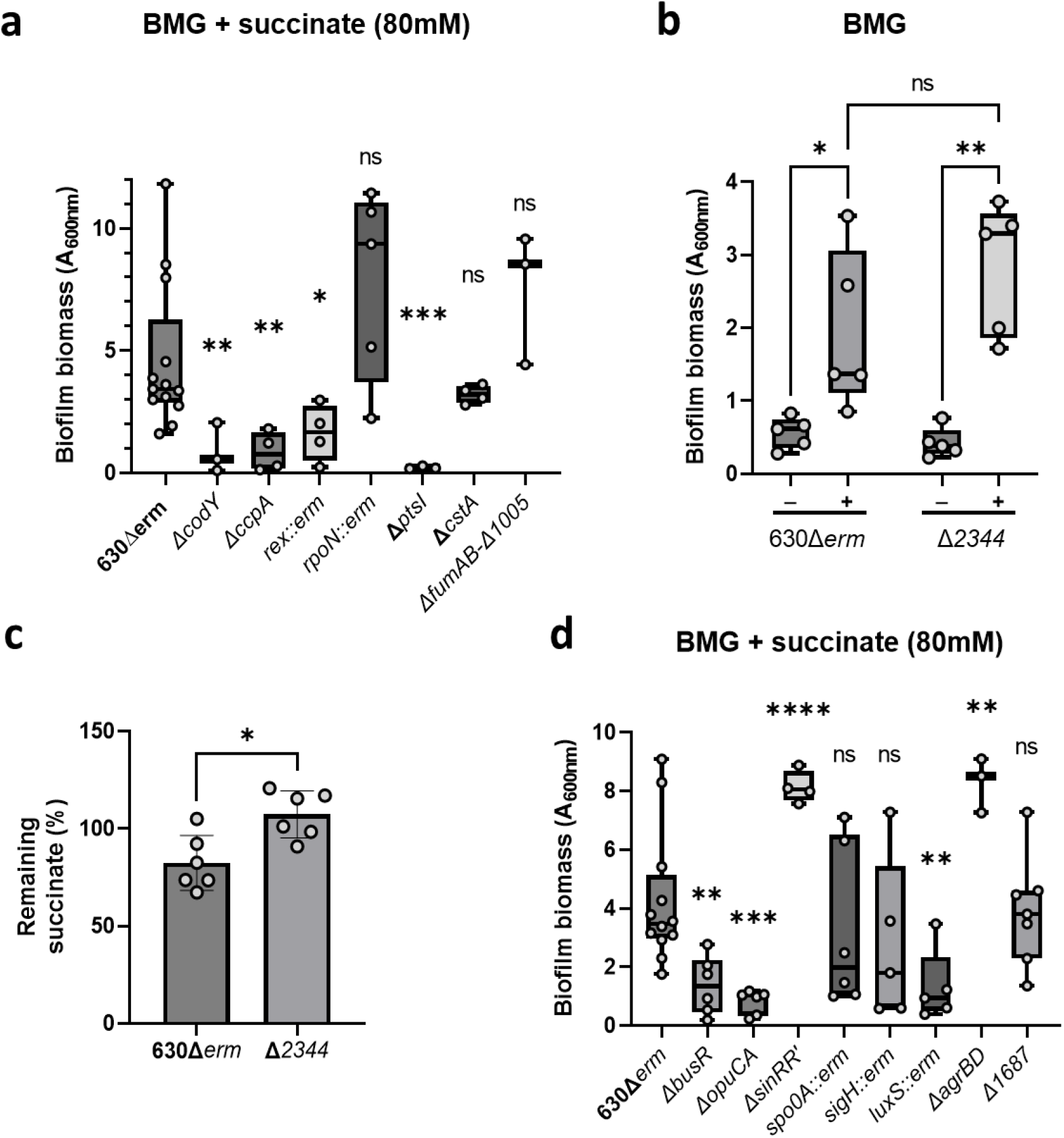
Extracellular succinate induces biofilm formation through global transcriptomic regulators. Biofilm formation was assayed in BMG medium supplemented with succinate (80mM) of **a**. various *C. difficile* 630Δ*erm* mutants in global regulations of metabolism (*ΔcodY, ΔccpA, rex::erm, rpoN::erm, ΔptsI, ΔcstA, ΔfumAB-Δ1005* and **b**. in the mutant for succinate intake *Δ2344* compared to the wild type strain. **c**. Measure of the remaining succinate in spent medium from cultures of the 630Δ*erm* and Δ*2344* strains in BMG medium supplemented with succinate (80mM) after 24h of incubation. **d**. Biofilm formation was assayed in BMG medium supplemented with succinate (80mM) of various *C. difficile* 630Δ*erm* mutants in osmotic stress response (*ΔbusR, ΔopuCA*), phase-related regulations (*ΔsinRR’, spo0A::erm, sigH::erm, luxS::erm, ΔagrBD, sigH::erm*), and *Δ1687* were tested and compared to the wild type strain. Statistical analyses performed are a,d) an ordinary one-way ANOVA test followed by an uncorrected Fisher’s LSD b) a Brown-Forsythe and Welch ANOVA test followed by an unpaired t-test with Welch’s correction and c) a paired t-test (ns: not significant; *: p<0.05; **: p<0.01; ***: p<0.001; ****: p<0.0001).

Finally, in contrast with 14h, many genes encoding cell wall components are up-regulated at 24h. Additionally to genes encoding peptidoglycan synthesis, they include genes of known surface proteins like fibronectin-binding protein A required for biofilm formation in *Staphylococci* (42,43) and cell wall proteins such as Cwp84, which modulates biofilm formation in *C. difficile* (44). In addition, several genes encoding putative membrane and cell wall proteins of unknown function as well as lipoproteins are up-regulated in the presence of succinate (Table S2). We noted that the lipoprotein CD1687 recently demonstrated to be essential in the DCA-induced biofilm (10,45) is not regulated by succinate and confirmed that a *CD1687* mutant is not affected in biofilm formation in the presence of this inducer (Figure 4a).

### Extracellular succinate induces biofilm formation through specific regulatory pathways

Since several known regulators involved in cell metabolism are differentially regulated in the succinate-induced biofilm, we tested the ability of mutants for metabolic regulators to form biofilm in response to succinate (Figure 4a). We showed that gene inactivation of the global metabolic regulators CcpA and CodY(34) strongly decreases biofilm formation in the presence of succinate. In addition,the *busR* and *opuCA* mutants are affected in their ability to form biofilm (Figure 4a). Thus, both carbon and nitrogen metabolism as well as osmotic stress response seem essential for succinate induced biofilm formation. The absence of genes of Strickland fermentation up-regulated and the down-regulation of *rpoN* suggests that Stickland reactions are probably not essential in succinate-induced biofilm formation. Interestingly, we observed that the *cstA* gene was up-regulated in the presence of succinate at 24h (Table S2). CstA is a pyruvate importer, which in absence of glucose is essential for pyruvate-induced biofilm formation (30). However, the *cstA* mutant did not display an altered biofilm response to succinate compared to the wild type 630Δ*erm* strain (Figure 4a). This indicated that the mechanism involving extracellular pyruvate as a key component for DCA-induced biofilm formation (30) is not the same for succinate induction.

Among the few genes up-regulated at both 14 and 24h, we found genes of the succinate utilization operon significantly up-regulated (table S2). Therefore, we deleted the *CD2344* gene encoding the succinate importer known to be essential in succinate utilization by *C. difficile* (22), and tested it for biofilm formation in the presence of succinate (Figure 4b). In parallel, we measured the concentration of succinate in the medium after 24h of incubation in the wild-type and mutant strains. While succinate was consumed by the wild-type strain at 24h, succinate concentration did not vary for the mutant strain, confirming that succinate intake is indeed impaired in the Δ*2344* mutant strain (Figure 4c). However, even as the Δ*2344* mutant strain is unable to import succinate, it is still able to form biofilm in response to this molecule (Figure 4b), suggesting it is the presence of extracellular succinate and not its consumption by *C. difficile* that would likely induce biofilm formation.

### Succinate induces the formation of a thick porous biofilm

To compare the morphology of *C. difficile* biofilms induced by succinate with that of DCA used as a reference, we performed confocal laser scanning microscopy (CLSM) analyses. We used SYTO9 and propidium iodide to detect live/dead cells, respectively. As shown in Figure 5a and Figure 5d, biofilms formed in the absence of any inducer had a larger dead population (red) than those induced by succinate or DCA. The latter seems to be much more abundant in living cells (green cells) and differently distributed according to the inducer. In addition, the shape, thickness, and global aspect of the biofilms are also highly dependent on the inducer. Indeed, succinate-induced biofilms are significantly thicker than DCA-induced biofilms and non-induced biofilms at 48h of incubation (Figure 5b). Moreover, succinate-induced biofilms display more complex shapes with a higher elevation than non-induced biofilms and DCA-induced biofilms, which have a flatter topography. These shapes are correlated with the spatial distribution of the dead cells, which is widely different between the inducers (Figure 5a). Indeed, while DCA-induced biofilms and non-induced biofilms have evenly distributed dead cells across the biofilm, succinate-induced biofilms present clusters of dead cells distributed throughout the biofilm mixed with the living cells. Succinate-induced biofilms also present a porous structure, with more space between the cells than non-induced and DCA-induced biofilms. This looseness can be measured with the roughness parameter (Figure 5c), calculated from an index based on the quantity of empty space between detected cells. For succinate-induced biofilms, the roughness is high at both 24h and 48h, while that of DCA-induced stays at a level comparable to the non-induced biofilm (Figure 5a, 5c). On the other hand, the biovolume of dead cells is lower in DCA and succinate-induced biofilms when compared to non-induced biofilms (Figure 5d). This is consistent with the cell survival of *C. difficile* already observed in the DCA-induced biofilm (Dubois *et al*. 2019). We concluded that succinate-induced biofilms appear more structured than DCA-induced and non-induced biofilms. This may be related to the distribution of dead cells within the extracellular matrix of the biofilm, inducing mechanical forces responsible for various structures, as already described for the biofilm structure of *B. subtilis* (46).

**Figure 5:**
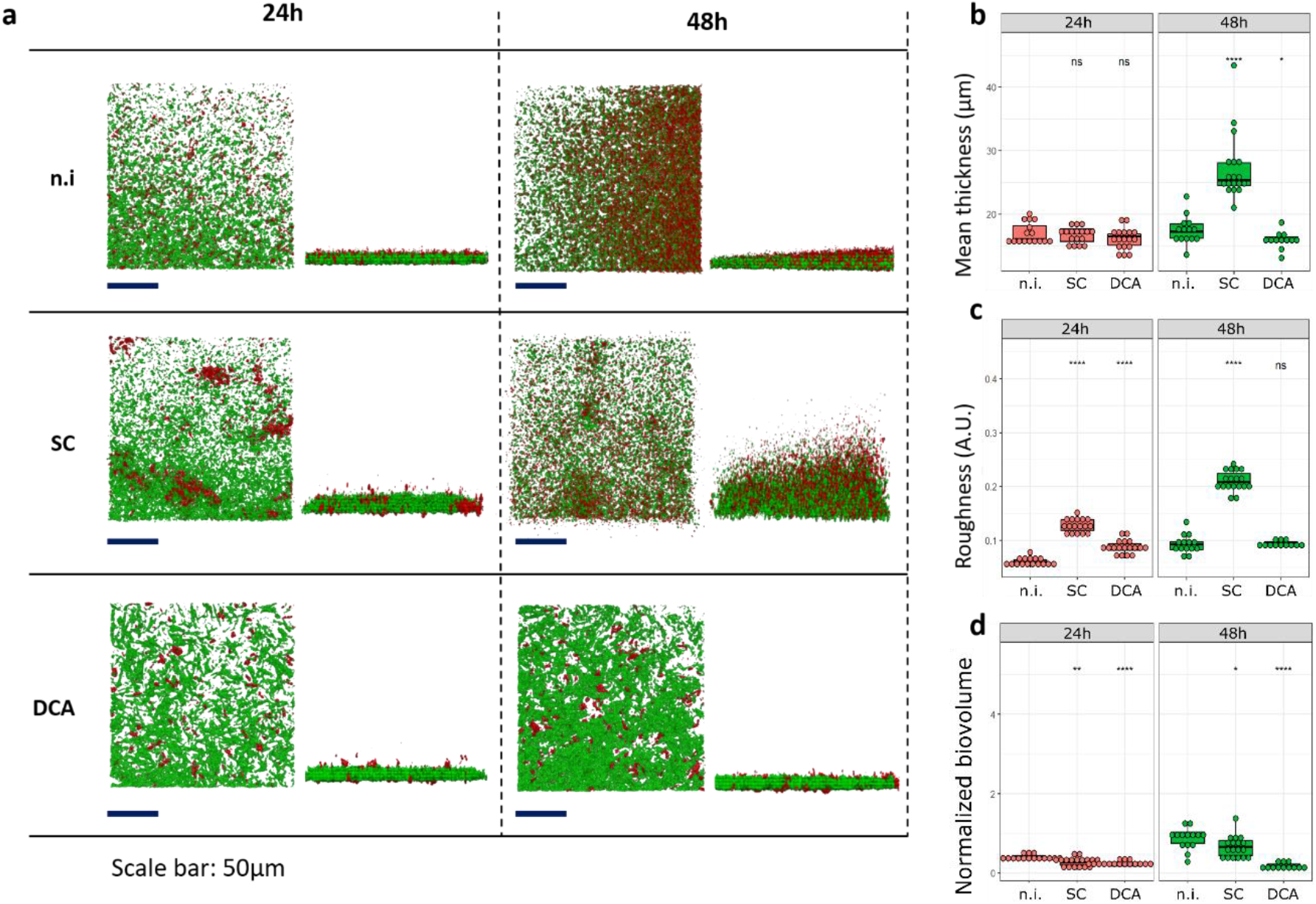
Live/dead observations of biofilms by CLSM. **a**. CLSM observation of 24h and 48h biofilms of the 630Δ*erm* strain grown in BHISG or BHISG media supplemented with either DCA (240μM) or succinate (120mM). Live cells are stained with Syto9 (green) and dead cells are stained with propidium iodide (red). Z-stacks were analysed with BiofilmQ and rendered with Paraview. Images are representative of 3 independent biological replicates. Quantitative analyses were performed with BiofilmQ to measure in **b**. the mean thickness of the biofilm of each image (measured with the SYTO9 signal), in **c**. the roughness of the biofilm (measuring space between live cells, with the SYTO9 signal) and in **d**. the normalized biovolume of dead cells (measured with the ratio of propidium iodide signal over SYTO9 signal). Each data point represents one technical replicate taken from three independent biological replicates. n.i.: no inducer; SC: succinate (120mM); DCA: deoxycholate (240μM). Statistical analyses performed here are Wilcoxon tests (ns: not significant; *: p<0.05; **: p<0.01; ****: p<0.0001). Scale bar: 50μm.

### Matrix composition of succinate-induced biofilms

Biofilm matrix is generally made of extracellular DNA (eDNA), polysaccharides, and proteins that hold cells together. In order to characterize the composition of the extracellular matrix of succinate-induced biofilms, we performed *in situ* CLSM analysis using fluorescent markers for cells and matrix components of succinate-induced biofilms, which we compared with CLSM analysis of non-induced and DCA-induced biofilms (Figures 6 and Figure S1). *C. difficile* cells of the 24h and 48h biofilms were stained either with Syto9 or Syto61 according to the matrix marker used to localize eDNA (TOTO-1), proteins (Sypro Ruby) and exopolysaccharides (EPS) such as β1-3 and β1-4 polysaccharides (calcofluor white). At 24h, extracellular proteins, eDNA and EPS (in red) of the non-induced and DCA-induced biofilms seem to be uniformly distributed among the cells (in green), while they are patchily distributed in succinate-induced biofilms (Figure 6a). We noted that the distributions of eDNA and proteins seem to follow similar patterns as the dead cells imaged with propidium iodide (Figures 5a and 6a), suggesting that eDNA and extracellular proteins could originate from dead cells. On the contrary, EPS appear uniformly distributed at the biofilm surface independently from the growth conditions. We noted that the matrix of succinate-induced biofilms is thicker compared to the non-induced or DCA-induced biofilms (Figure 6b), this thickness does not seem to depend on the considered polymer. On the other hand, the normalized biovolume of each matrix component depends on the inducer, as well as on the considered polymer (Figure 6c). Although there is greater variability in eDNA production in succinate-induced biofilms, this component quantity does not seem to be affected by the inducer. On the opposite, succinate-induced biofilms produce less extracellular proteins and EPS than non-induced and DCA-induced biofilms. Thus, as already shown by Dubois and collaborators (2019), we confirmed in all biofilm conditions that proteins and eDNA are the main components of the matrix for the DCA-induced biofilm.

**Figure 6:**
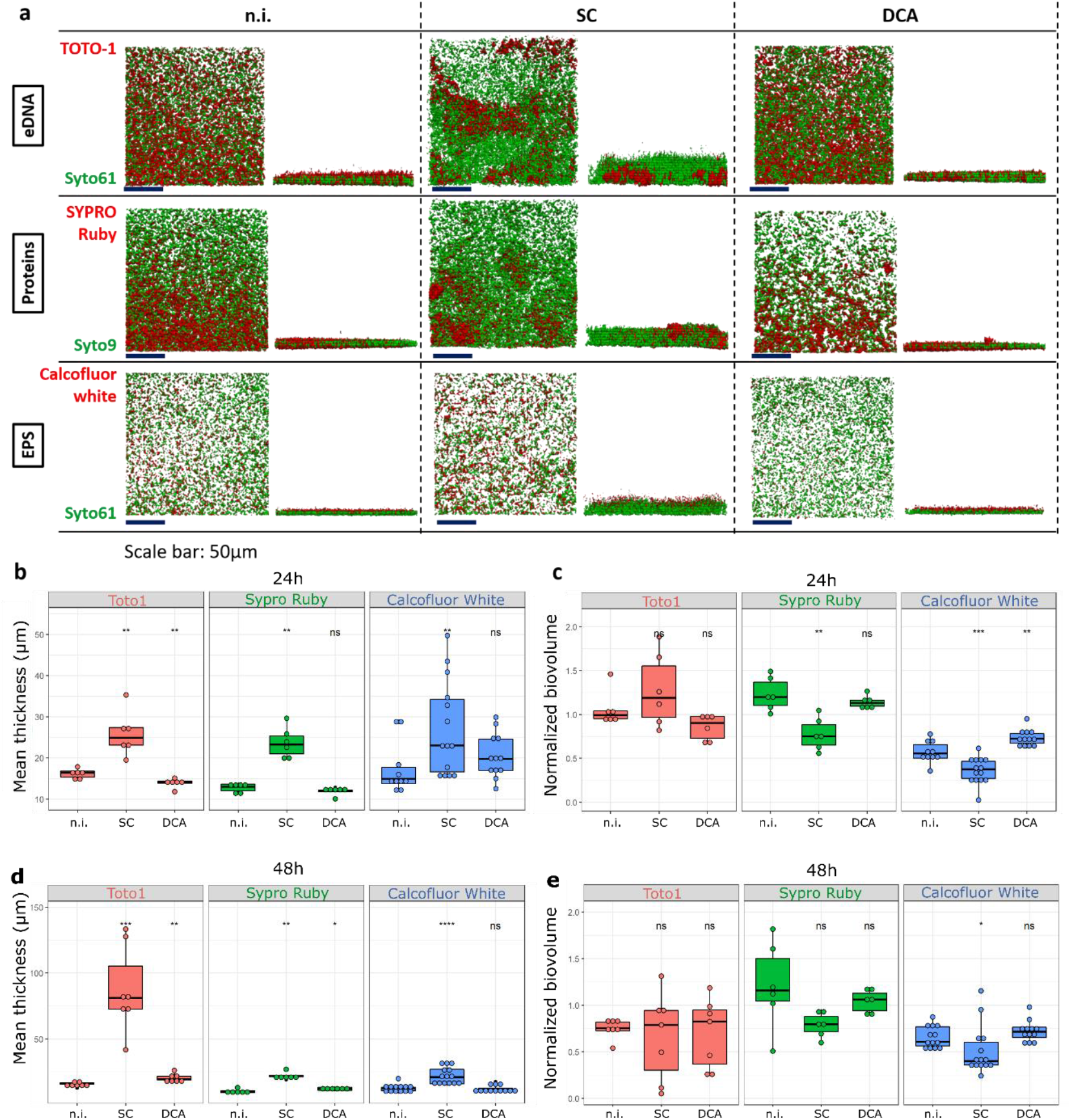
CLSM observations of biofilm matrix at 24h of incubation. **a**. CLSM observations of 24h biofilms of the 630Δ*erm* strain grown in BHISG or BHISG media supplemented with either DCA (240μM) or succinate (120mM). Biofilm matrix components were marked with either TOTO-1 (eDNA), Sypro Ruby (proteins) or calcofluor white (β1-3 and β1-4 polysaccharides), and they appear in red. Cells were marked either with Syto9 or Syto61, depending on the other marker used and they appear in green. Z-stacks were analysed with BiofilmQ and rendered with Paraview. Images are representative of 3 independent biological replicates. Quantitative analyses were performed with BiofilmQ to measure in **b**. and **d**. the mean thickness of the observed matrix components of each image at 24h and 48h, respectively, and in **c**. and **e**. the normalized biovolume of the observed matrix components (measured with the ratio of the biovolume of the considered component over the biovolume of relevant SYTO signal). Each data point represents one technical replicate taken from three independent biological replicates. n.i.: no inducer; SC: succinate (120mM); DCA: deoxycholate (240μM). Statistical analyses performed here are Wilcoxon tests (ns: not significant; *: p<0.05; **: p<0.01; ****: p<0.0001). Scale bar: 50μm.

At 48h of incubation, EPS are also uniformly distributed at the top of the cells for all conditions and are less predominant than eDNA and proteins (Figure S1). In these later biofilms, proteins seem to be clustered in all conditions, although to a lesser extent in the succinate-induced biofilm, which seems not to be linked to dead cells as they do not follow patterns of the dead cells imaged (Figure 5a). Surprisingly, an important quantity of eDNA is localized at the cell surface and spreads quite far above the succinate-induced biofilm (Figure 6d and Figure S1), although its biovolume is not statistically different from the other conditions (Figure 6e). The differences between the growth conditions in normalized biovolume of matrix components seem to lessen at 48h (Figure 6e) even though there are still less extracellular proteins and EPS in succinate-induced biofilms. Overall, when induced by succinate, *C. difficile* biofilms are thicker and less protein- and EPS-rich than the other described biofilms, and the distribution of eDNA is widely different from the other conditions. Thus, different inducers can promote the formation of *C. difficile* biofilms, which may differ in structure and characteristics.

## Discussion

Succinate present in the dysbiotic intestine in relatively high concentrations is associated to the regulation of the immune response against bacterial infections, and it is also involved in energy metabolism in *C. difficile* (22,47). Therefore, it was reasonable to assume that succinate could induce biofilm formation through direct modulation of metabolism. However, we showed that a succinate importer (CD2344) mutant was still able to produce biofilm in the presence of succinate, suggesting that extracellular succinate, and not its use in metabolism, induces biofilm formation. This is consistent with the capacity of the *fumAB-CD1005* mutant to produce biofilm when succinate is added to the growth medium (Figure 4a). The ways by which *C. difficile* can sense extracellular succinate is an open question. In *Bacillus subtilis*, the DctABS system forms a functional sensor for C4-dicarboxylates such as succinate, fumarate or malate and transduces the signal to the DctR response regulator (48). Unfortunately, only one *C. difficile* strain (NCTC13750) displays a protein homologous to DctS and DctA. The candidates for succinate sensing in *C. difficile* could be found among the eight up-regulated two-component histidine kinase of unknown function detected in our RNAseq experiments (Table S2). In addition, extracellular succinate may be sensed through cell-wall proteins or membrane modifications. This has been described in *B. subtilis* with the protein KinC, a membrane histidine kinase that detects potassium leakage caused by the action of surfactin from across the membrane (49). Then the signal is transmitted through the phosphorylation of the response regulator Spo0A, which in turn controls the activity of the antirepressor SinR leading to biofilm formation and other cell mechanisms including sporulation (50,51). In *C. difficile* such a scenario may involve the regulatory response CsfT protein, upregulated in response to succinate both at 14h and at 24h (Table S2). CsfT is an ECF sigma factor associated with signal transduction of extracellular signals such as cell wall modifications through lysozyme (38). This sigma factor could therefore be directly associated with the transmission of the extracellular succinate signal in case of cell wall modifications. On the other hand, lipoproteins which are involved among other things in nutrient uptake and signalling systems (52), may be able to detect extracellular succinate. We recently demonstrated that the lipoprotein CD1687 of the 630Δ*erm* strain mediates in part the metabolic reorganization occurring in DCA-induced biofilm (10,45). Since CD1687 is not involved in succinate-induced biofilm formation, we cannot exclude that lipoproteins up-regulated in the presence of succinate (Table S2), could sense succinate and/or induce metabolic pathways involved in biofilm formation. Deletion mutants for their encoding genes will be needed to verify whether these candidates are involved in succinate detection and biofilm formation.

We showed that succinate is a strong biofilm inducer for several *C. difficile* strains, and compared with the DCA-induced biofilms, the succinate-induced biofilms are thicker. This increased thickness seems associated with a more complex 3D structure organized around cell death distribution, releasing DNA and cellular content into the environment after 24h of incubation. In *B. subtilis* and *Pseudomonas aeruginosa*, such complex structures are responsible for wrinkles on the biofilm surface (53,54) and are likely due to localized cell death (46,55). Similar structures have been observed in *C. difficile* (8). We also noted that at 48h of incubation, proteins form clusters in all conditions, although this should be interpreted cautiously as PFA is known to form such protein aggregates. Interestingly, we observed that EPS are uniformly localized on top of the cells for all conditions tested, which is consistent with the expression of the *ccsA* gene (*CD630_25450*) encoding *C. difficile*’s cellulose synthase (56), not significantly differentially regulated (Table S2) (10). Such distribution of EPS is probably due to their production only by cells localized at the top of the biofilm, by a division of labor similar to what was observed in *B. subtilis* (57). The most striking feature of the succinate-induced biofilms is the mass of eDNA above the cells. As the biovolume of eDNA detected in succinate-induced biofilms is similar to that of the DCA-induced biofilms (Figure 6, Figure S1), such a mass of eDNA is unlikely to participate in the structure of biofilms formed in the presence of succinate. On the other hand, the decrease in protein production observed in the succinate-induced biofilm could partly explain the more porous structure of succinate-induced biofilms.

With over half of the genome differentially regulated at 24h, the transcriptional state of the cells present in the succinate-induced biofilm is strongly modified compared to planktonic cells. In biofilms induced by succinate, cells are more prone to divide and grow, as seen by the up-regulation of genes involved in cell-wall synthesis, division, fatty acid biosynthesis and importation of sugars likely used in cell-wall synthesis. Metabolism genes are also affected and mainly shift from glycolysis, Wood-Ljungdahl and Stickland fermentation pathways to the utilization of succinate. In other conditions of the *C. difficile* biofilm formation, the Stickland fermentation pathways were up-regulated compared to the planktonic cells, especially the utilization of branched-chain amino-acids (29,30). Thus, this metabolic pathway does not seem essential for biofilm formation in response to succinate, which is consistent with the down-regulation of the *rpoN/sigL* gene expression at 24h of the succinate-induced biofilm, known to regulate the expression of the Stickland fermentation genes (58). Thus, biofilm formation in the presence of succinate is associated in part with metabolic rearrangements, as already observed with deoxycholate (30). However, the modifications to the metabolic pathways are not strictly the same, suggesting that biofilms can be promoted by different mechanisms of cell adaptation.

While they share similarities with other gut metabolite-induced biofilms, biofilms induced by succinate differ in several aspects. First, we observed a more complex spatial organization of succinate-induced biofilms than non-induced or DCA-induced biofilms, which appear strongly associated with a large distribution of the live/cell death in such an environment. Secondly, the circumstances of their formation are quite different from those of most biofilms already described, and those are mainly formed in response to stressful conditions due to the presence of metronidazole (9) or DCA (10) or during the late stationary phase (59). In fact, biofilm induction in response to succinate appears to occur through the signalling of a key energy metabolite such as NAD+ whose regeneration is primarily associated with the use of succinate. In agreement, succinate seems more favorable to *C. difficile*’s growth than the other inducers such as deoxycholate, which significantly affects the growth rate of *C. difficile* when added to the medium (60). However, we cannot exclude that succinate-induced biofilms can be mediated through osmotic regulation suggested by the expression of *busR*-like and *opuCA* genes as well as the synthesis and importation of the osmoprotectant ornithine, spermidine, and ornithine involved in the osmotolerance of *C. difficile* (41).

Finally, the importance of succinate as an inducer of *C. difficile* biofilm formation must be taken into account in relation to its availability in the gut. Indeed, in healthy gut microbiota, succinate is produced and consumed by commensal bacteria, achieving sub-millimolar to millimolar concentrations of luminal succinate (47,61). However, in a dysbiotic microbiome depleted of its succinate consumers, the mean succinate concentration is measured in tens of millimolar, which are the concentrations used in this study (47,61) and encountered by *C. difficile* during CDI after intestinal dysbiosis. Thus, elevated concentrations of succinate in the gut can be viewed both as a nutrient source (22) and as an inducer for biofilms, independently of its metabolism, as shown in this study. Biofilms are known to increase resistance to treatments therefore, understanding the mechanisms of biofilm formation by *C. difficile* and their characteristics becomes urgent in order to adapt treatment strategies and take potential biofilms into account.

## Acknowledgments

Pierre Adenot and Vlad Costache from the INRAE MIMA2 imaging platform are acknowledged for their assistance with CLSM and Axel Ranson from Laboratoire Pathogenèse des Bactéries Anaérobies and Ecole Normale Supérieure Paris-Saclay for his help in biofilm experiments. This work was funded by the Institut Pasteur, the “Integrative Biology of Emerging Infectious Diseases” (LabEX IBEID) funded in the framework of the French Government’s “Programme Investissements d’Avenir” and The ANR DifBioRel AAPCE5. EA is a doctoral fellow of Université Paris-Cité.

