## Supplementary Figure S1 for "Extracellular succinate induces spatially organized biofilm formation in *Clostridioides difficile*"

### Supplementary material

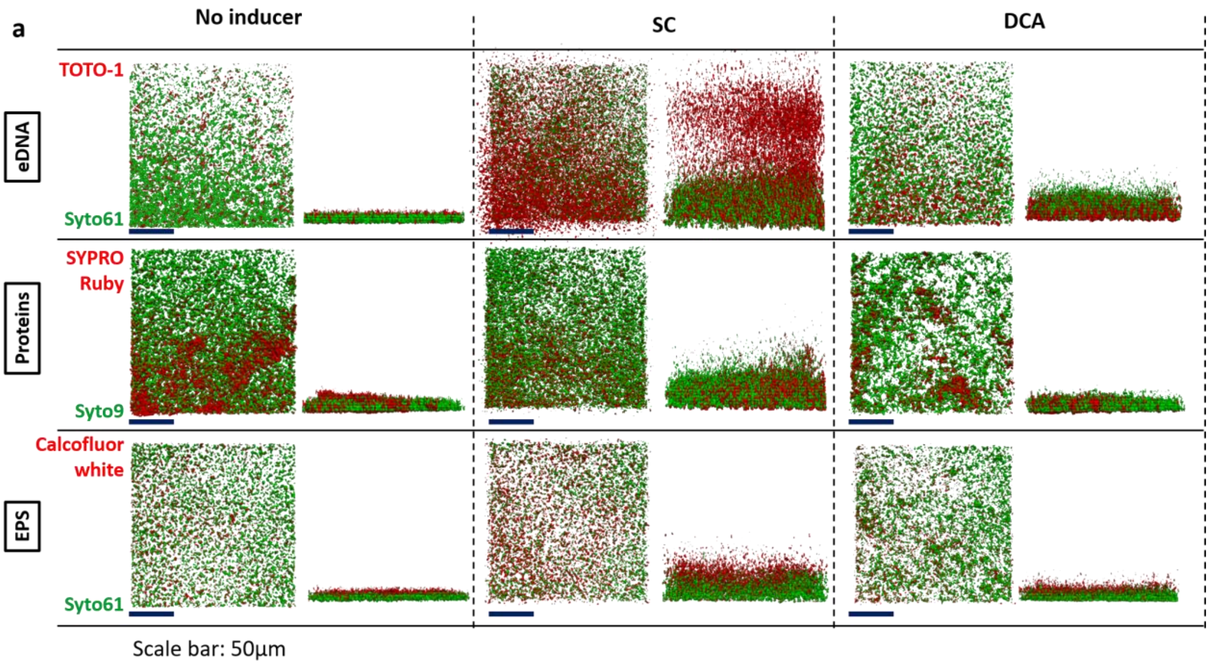


**Figure S1: CLSM observations of biofilm matrix at 48h of incubation.**
CLSM observations of 48h biofilms of the 630Δ*erm* strain grown in BHISG or BHISG media supplemented with either DCA (240µM) of succinate (120mM). Biofilm matrix components were marked with either TOTO-1 (eDNA), Sypro Ruby (proteins) or calcofluor white (β1-3 and β1-4 polysaccharides), and they appear in red. Cells were marked either with Syto9 or Syto61, depending on the other marker used, and they appear in green. Z-stacks were analyzed with BiofilmQ and rendered with Paraview. Images are representative of 3 independent biological replicates. Scale bar: 50µm. n.i.: no inducer; SC: succinate (120mM); DCA: deoxycholate (240µM).
